# Structural basis of phosphate export by human XPR1

**DOI:** 10.1101/2024.08.22.609128

**Authors:** Qixian He, Ran Zhang, Sandrine Tury, Valérie Courgnaud, Fenglian Liu, Jeanluc Battini, Baobin Li, Qingfeng Chen

## Abstract

Phosphorus is an essential element for all living organisms. While inorganic phosphate (Pi) cellular import is well documented, export is poorly characterized despite the recent discovery of the inositol pyrophosphate (PP-IP)-dependent Pi exporter XPR1. In this study, we determined the cryo-EM structures of XPR1. XPR1 forms a loose dimer, with each protomer containing 10 transmembrane helices (TMs) that can be divided into a peripheral domain (TM1, 3 and 4) and a core domain (TM2 and TM5-10) structurally related to ion-translocating rhodopsins. Bound Pi is observed in a tunnel formed by the α helical bundle inside the core domain at a narrowest point that separates the tunnel into an intracellular vestibule (IV) and an extracellular vestibule (EV). This narrowest point contains a cluster of Pi-coordinating basic residues, and their substitution by mutagenesis strongly impaired phosphate export. PP-Ips stimulate XPR1 activity by binding to its SPX domain. Loss of this interaction induces a conformational change of the wide open EV (conducting state) into a collapsed EV (closed state), mainly caused by structural movements of TM9 and the bulky sidechain of Trp573 right above the Pi binding site, which are essential for the Pi export activity of XPR1. Our structural and functional characterization paves the way for further in-depth studies of XPR1.

## Introduction

Phosphorus is an element that is essential for almost all biological processes^1^, but is toxic to cells in excess^2^. Therefore, living organisms have evolved various protein machines to maintain the phosphate homeostasis in cells. In human cells, multiple solute transporter (SLC) families are responsible for uptake of phosphate into cells, including SLC20 and SLC34 family, mostly in the form of inorganic phosphate (Pi)^1^. Na-phosphate cotransporter SLC34A2 (NaPi-IIb) is involved in active uptake of dietary Pi in the small intestine under Pi-limiting condition^3^. Absorbed Pi are mostly stored in bones, and can be mobilized under low-Pi condition, a process regulated by parathyroid hormone (PTH) and 1,25(OH)_2_ vitamin D_3_^4^. Circulating Pi are filtered in glomeruli, and most of the filtered Pi are reabsorbed in the proximal tubular epithelial cells and the distal tubule by SLC34A1 (NaPi-IIa) and SLC34A3 (NaPi-IIc), which is tightly regulated by PTH and fibroblast growth factor 23 (FGF-23) via controlling the availability of these transporters^5^. By contrast, members of SLC20 are ubiquitously expressed in all cells and responsible for providing Pi for normal cellular structure and function^6^.

Compared with abundance of studies on proteins involved in Pi uptake, less is known about the proteins involved Pi export. Xenotropic and Polytropic Retrovirus Receptor 1 (XPR1), or SLC53A1, which was initially identified as a receptor for xenotropic and polytropic murine leukemia retroviruses (X- and P-MLV) ^7–9^, has been shown to function as a Pi exporter^10,11^, and a key regulator of cellular Pi homeostasis^12^. Harboring a SPX (SYG1/Pho81/XPR1) domain which is capable of sensing inositol polyphosphate^13^ and a transmembrane domain with multiple transmembrane helices, XPR1 mediates transcellular Pi export in an inositol pyrophosphate (PP-IP)^12,14,15^. Knockout of diphosphoinositol pentakisphosphate kinases (PPIP5Ks) or inositol hexakisphosphate kinases (IP6Ks), which are involved in synthesis of PP-IP, or pharmacological inhibition of IP6Ks, resulted in inhibition of XPR1-mediated Pi efflux^12,14,15^. Interestingly, the envelope-receptor-binding domain (XRBD) of X-MLV was shown to be able to inhibit XPR1-mediated Pi efflux^10^.

XPR1 is the subject of growing interests as it is related to several human diseases presenting Pi homeostasis dysfunction. Indeed, 1) XPR1 variants are found in patients with primary familial brain calcification (PFBC)^16–20^, a neurodegerative disease characterized by calcium phosphate deposits in cerebral microvessels due to dysfunction of cellular Pi homeostasis^21^; 2) XPR1 was shown to modulate intracellular polyphosphate content in platelet^22^, and its inactivation resulted in polyphosphate accumulation, accelerated arterial thrombosis, and augmented activated platelet-driven pulmonary embolism in mice^22^; 3) XPR1 promotes the progression and is a diagnostic marker of tongue squamous cell carcinoma^22–24^; 4) *XPR1* has been identified as a therapeutic vulnerability gene in ovarian and uterine cancers, as inhibition of XPR1-mediated phosphate efflux reduced tumor cell viability through toxic accumulation of intracellular phosphate^25^. Thus, XPR1 represents an interesting target for drug development for intervention in diseases with compromised Pi metabolism.

Despite several studies have suggested that XPR1 is a Pi exporter, its Pi export activity has been questioned due to the lack of definitive evidence such as the three-dimensional structure of XPR1 and structure-function studies^10–12,14,26,27^. To fill this gap, we determined cryo electron microscopy (cryo-EM) structures of human XPR1 in two distinct conformations, which reveal that it forms a dimer, and that the transmembrane core domain of XPR1 harbors a Pi binding site and a translocation pathway structurally similar to ion-translocating microbial rhodopsins. Moreover, two conformations are likely in conductive and non-conductive state respectively. These structural observations, which are complemented with site-directed mutagenesis, Pi transport assay, and molecular dynamic (MD) simulations, confirmed the functional role of XPR1 as a Pi exporter, revealed its Pi translocating pathway, and provided a framework for further mechanistic studies on this family of proteins. Our studies also pave the way for structure-based drug design targeting XPR1.

## Results

### Structure determination and overall structure

Topological wise, XPR1 is known to contain an N-terminal SPX domain, a transmembrane domain (TMD) and a C-terminal domain (CTD). While the TMD appears to be responsible for Pi translocation across cell membrane, the intracellular SPX domain was reported to be involved in InsP_8_ binding, which is an inositol pyrophosphate of endogenous source and essential for phosphate translocation activity of XPR1^14^. Full length XPR1 fused a C-terminal Strep tag (referred to as XPR1^wt^ hereafter) were expressed in HEK293F cells. After purification, XPR1^wt^ was subjected to single particle cryo-EM analysis, which yielded an electron density map at an overall resolution of 2.9 Å (Supplementary Fig. 1). While the TMD is well resolved (Supplementary Fig. 2), the intracellular SPX domain is completely disordered, implying its high intrinsic flexibility (Fig. 1a-b).

**Figure 1.**
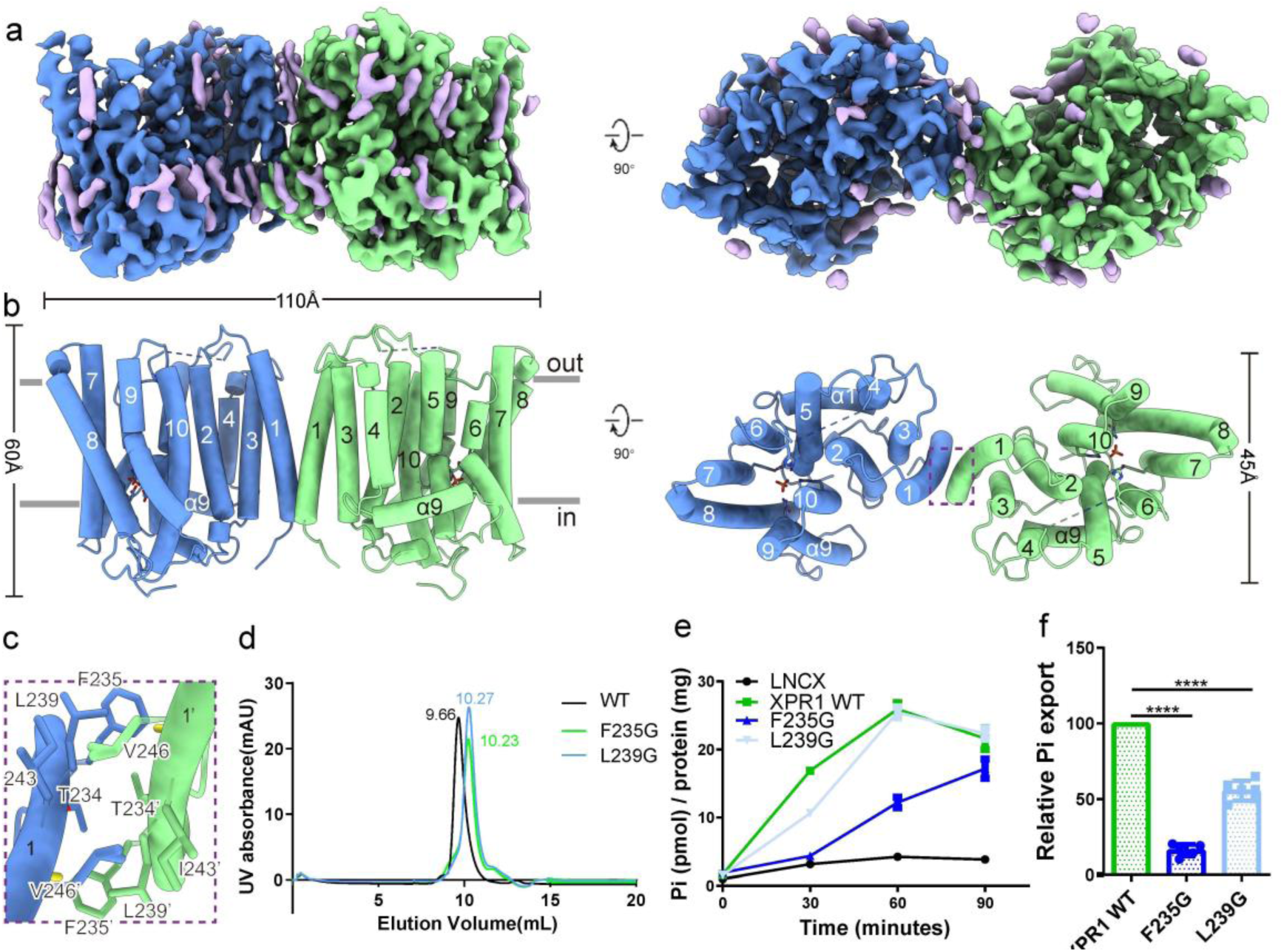
Overall structure of XPR1. **a-b**, Electron density maps **(a)** and cartoon representations **(b)** of XPR1 in two views. **c**, Expanded view of the dimer interface of XPR1 (the region boxed in **b**), highlighting key residues involved in dimer formation. **d**, Disruption of XPR1 dimer by mutation of single residues in the dimer interface as shown by SEC. **e**, Kinetics of Pi export in HCT116 XPR1 KO cells expressing XPR1 WT or dimer interface mutants (F235G, L239G). Pi released in culture medium was quantified by malachite green phosphate assay and normalized to the total amount of proteins. Data are means ± SD from a representative experiment (n=6). **f**, Relative Pi export level in cells expressing dimer interface mutants as compared with Pi export level measured in XPR1 WT cells at time point of 30 min. n=6 independent experiments. One-way ANOVA with Dunnett’s multiple comparisons test * = p ≤ 0.05, **** = p ≤ 0.0001.

XPR1 adopts an elongated homo-dimeric architecture, as manifested by the dimension of XPR1 dimer (110 Å x 60 Å x 45 Å) (Fig. 1b). There are 20 transmembrane helices (TMs) in total, with 10 TMs in each protomer (Fig. 1a-b). Surprisingly, only TM1 in each protomer participates in dimer interface formation, and two TM1s are tilted from each other, with minimal contacts formed only at their N-termini (Fig. 1b-c). Therefore, interactions in the dimer interface in TMD are very sparse (with a buried surface of ∼454.8 Å^2^) and only a few residues from TM1 of each protomer are involved, including Thr 234, Phe 235, Leu 239, Ile 243, and Val 246 (Fig. 1c). As for the C-termini of two TM1s, they are too far away from each other to form direct interactions, leaving a void in between (Fig. 1b). This void is occupied by lipid molecules, which probably contribute to stable dimer formation in XPR1, and strengthen the sparse interactions formed at the N-termini of TM1s (Fig. 1b). To test whether dimerization of XPR1 is essential for its Pi export activity, we mutated these residues to glycine, and found that F235G and L239G are in mostly monomeric form, as shown in size exclusion chromatography, cross-linking experiments and Western blotting (Fig. 1d and Supplementary Fig. 3).

Pi export assay was conducted in human HCT116 cells, in which the endogenous XPR1 was knocked out (KO) using CRISPR/cas9 technology, resulting in loss of XPR1 cell surface expression and impairment of Pi export (Supplementary Fig. 4). XPR1 variants as well as XPR1^wt^ were re-introduced into the XPR1 KO cells and tested for Pi export activity. We showed that reintroduction of XPR1^wt^ in XPR1 KO cells restored the Pi export defect seen in XPR1 KO cells with or without reintroduced empty vector (Fig. 1e and Supplementary Fig. 4c). Among the dimer interface mutants that resulted in monomeric transporters, L239G presented a slightly weaker activity when compared to WT, which correlated with a slight decrease in plasma membrane expression (Fig. 1f, Supplementary Fig. 5). Pi export activity of F235G was dramatically diminished, again correlating with a compromised cell surface expression (Fig. 1f, Supplementary Fig. 5). Therefore, F235G and L239G likely affected targeting of XPR1 to cell membrane, rather than its intrinsic Pi export activity. Taken together, our data suggest that dimer formation may not be essential for Pi export by XPR1. Whether dimerization is a prerequisite for cell surface expression of XPR1 remains to be investigated.

### Protomer structure

To find out proteins that show structure similarity to XPR1, we performed a DALI search^28^ against all protein with known structure. DALI search shows that, while structure of TM1, 3, 4 do not seem to show high similarity to other structures in the database, TM2 and TM5-TM10 in each protomer form a structure similar to bacterial Na^+^/Cl^-^ transporting rhodopsin or channelrhodopsin. Structural homology to these proteins strongly suggests that XPR1 might function as a Pi exporter, as these regions might form the ion translocation pathway for Pi export. We named the region spanning TM2 and TM5-TM10 as the core domain, whereas the region spanning TM1, 3, 4 was named as the peripheral domain (Fig. 2a). Intriguingly, a non-protein density that fits nicely to a lipid molecule is observed in each protomer, in a cavity formed at the interface between the core domain and the peripheral domain, with openings to the intracellular membrane surface and lipid bilayers (Fig. 2a). The bound lipid, which is tentatively modeled as POPC, adopts a pose with its head group pointing to the surface of inner leaflet of the membrane and its acryl chains buried deeply inside the membrane (Fig. 2a). Whether and how this lipid molecule participates in regulating the function of XPR1 await further investigations.

**Figure 2.**
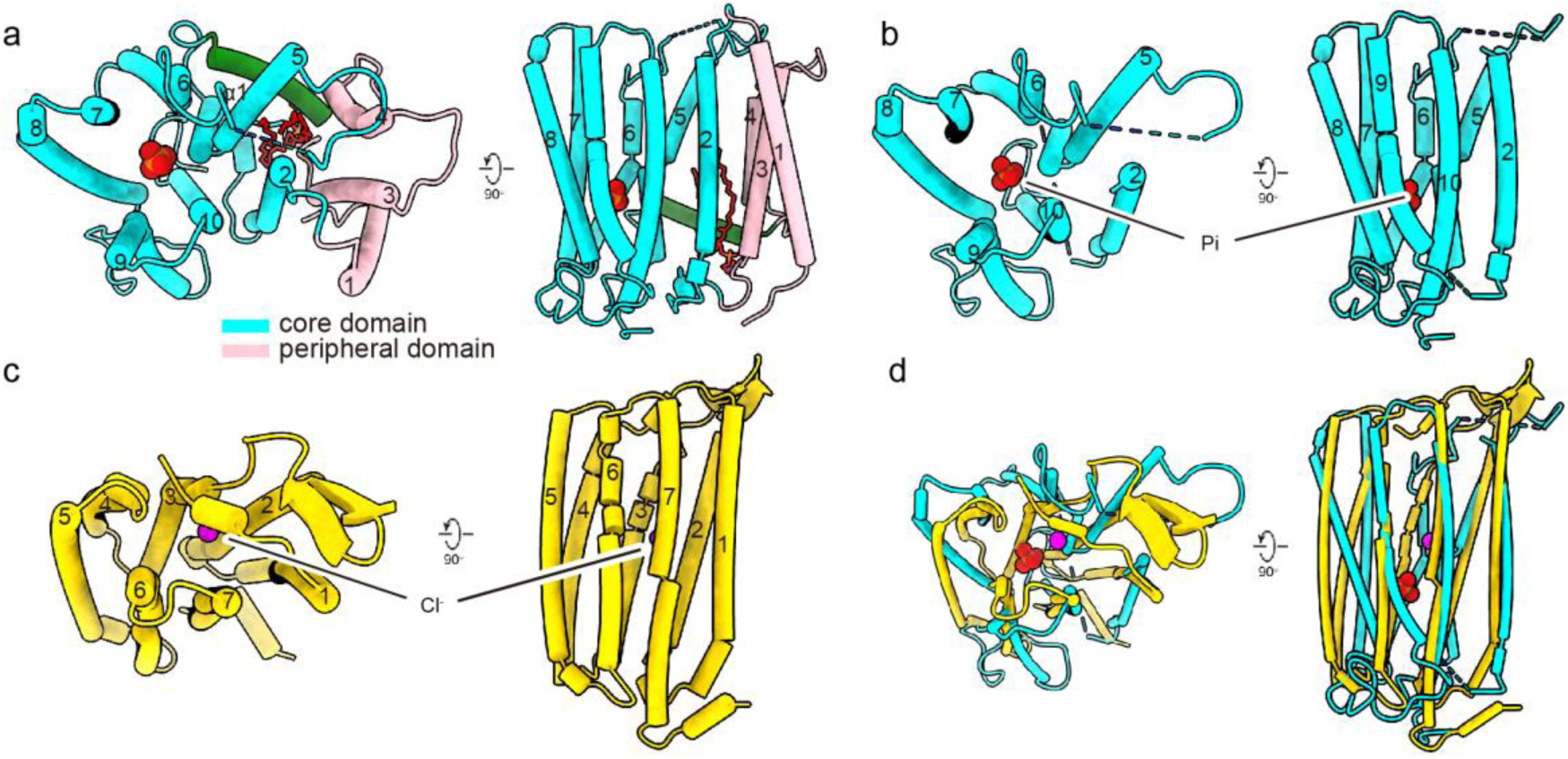
Protomer structure of XPR1 reveals structural similarity to ion-translocating rhodopsin. **a**, Protomer structure of XPR1 shown in two views, which can be separated into a core domain and a peripheral domain, with the core domain showing similarity to ion-translocating rhodopsin. Bound lipid is shown as red sticks. **b,** Structure of the core domain of XPR1 in two views. **c**, Structure of Cl^-^-translocating rhodopsin (pdb# 5b2n) protomer in two views. **d**, Overlay of structures of the core domain of XPR1 and Cl^-^-translocating rhodopsin in two views. Bound Pi in XPR1 and bound Cl^-^ in Cl^-^-translocating rhodopsin are shown as red and magenta spheres respectively.

Among proteins that are structurally similar to the core domain of XPR1 according to DALI search, chloride pumping rhodopsin (CIR) from *nonlabens marinus* (pdb# 5b2n) ^29^ is the topmost hit, albeit with only ∼8% sequence identity and an RMSD of ∼4 Å when their structures were superimposed. When viewed from the extracellular space, both of their 7-TMs (TM2 and TM5-10 in XPR1, and TM1-7 in rhodopsin) are arranged sequentially in counter-clockwise direction and form a helical bundle (Fig. 2b-d). Despite similar overall arrangement in TMs, there are striking structural differences. First, when compared with helices in rhodopsin, some of the helices in XPR1 are severely bent in the middle towards the center of ion translocation pathway, including TM5, 6 and TM9, which facilitate the formation of a narrowest point in the membrane for Pi coordination (Fig. 2b-d). As a result, the helical bundle formed in XPR1 appears to be more compact whereas the one formed in rhodopsin is more expanded (Fig. 2b-d). Second, similar to that of eukaryotic class A G-protein-coupled receptors, a C-terminal amphipathic helix is observed in CIR, which is parallel to and interacts with membrane surface, and may be important for the stability of CIR. On the contrary, C-terminus of XPR1 is largely disordered and a similar amphipathic helix is unlikely also present (Fig. 2b-d). Third, the ion binding site is positioned in the lower leaflet (closer to intracellular surface) and upper leaflet (closer to extracellular surface) of the membrane in XPR1 and CIR respectively (Fig. 2b-d). When viewed from above, Cl^-^ binding site in CIR is positioned roughly in the center of the 7-TM helical bundle, whereas in XPR1, its first TM (TM2) of the 7-TM helical bundle is not directly involved in Pi binding, therefore the position of Pi site deviates away from the center of the helical bundle (Fig. 2b-d).

### Pi translocation pathway and Pi binding site

An hourglass-shaped phosphate translocation pathway is located in the core domain of each XPR1 protomer, which is separated into an extracellular vestibule (EV) and an intracellular vestibule (IV) by a narrowest point in the middle of membrane (Fig. 3a-b). While all TMs in the core domain participate in forming the EV, only TM5-TM10 are involved in the formation of IV (Fig. 3a-b). Multiple layers of positive charged residues are observed along the phosphate translocation pathway (Fig. 3c), e.g., at the entrance of EV (Arg 270, Arg 273, Lys 442, Arg 448, and Lys 500), the entrance of IV (Arg 472, Arg 466, Arg 611 and Arg 624), and the narrowest point in the middle of the membrane (Arg 459, Arg570, Arg 603 and Arg 604), respectively. Among these positively charged residues, Arg 448 and Arg 459 have been identified in PFBC patients^18,19^. The narrowest point, positioned ∼17 Å (about 1/3 the thickness of lipid bilayer) away from the membrane surface at intracellular side, likely harbor the Pi binding site, which is formed by residues from TM5-TM10, but not TM2 of the core domain (Fig. 3c).

**Figure 3.**
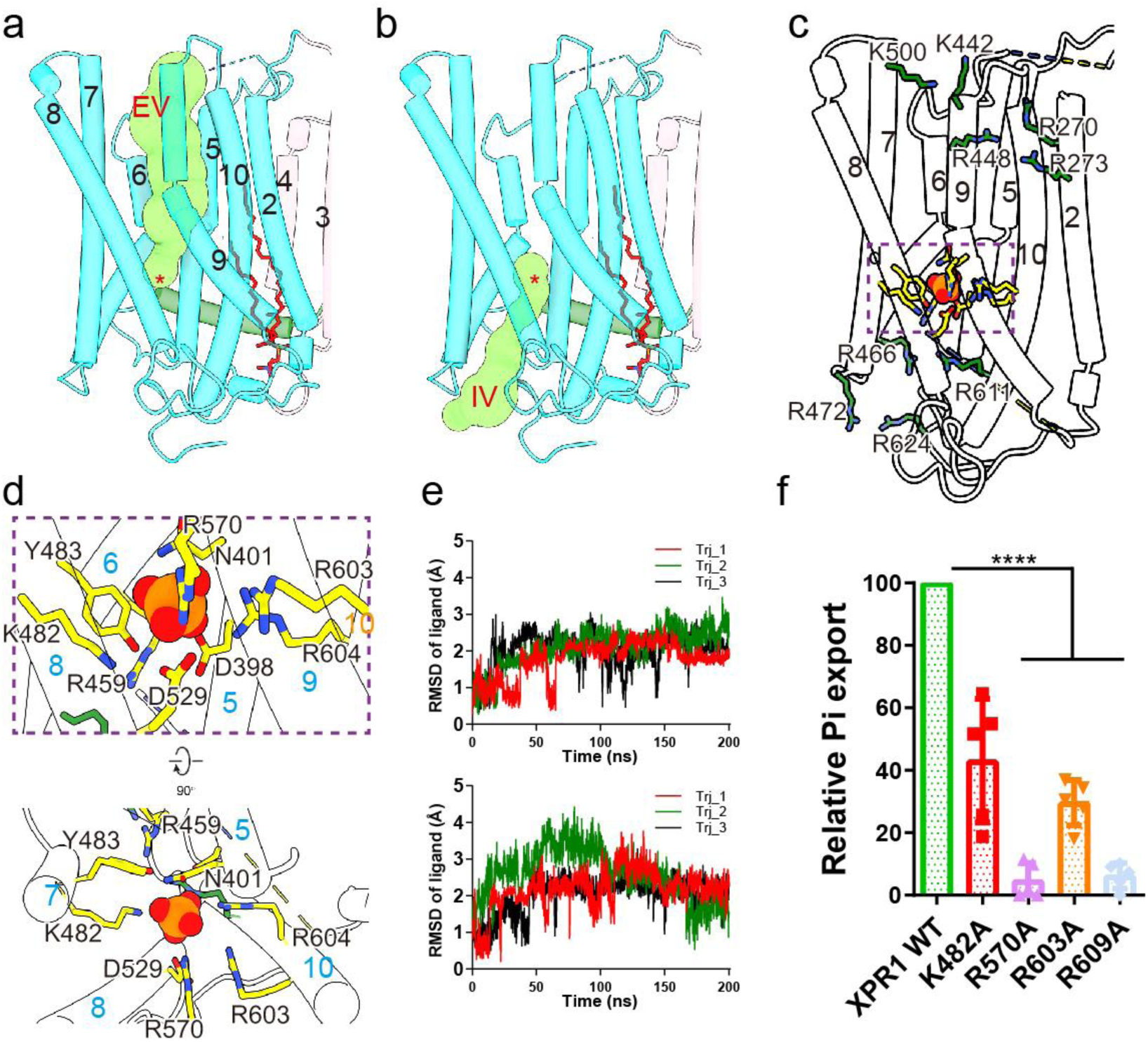
Pi translocation pathway and Pi binding site of XPR1. **a-b**, Pi translocation pathway in the core domain of XPR1, as calculated by Caver, showing EV (**a**) and IV (**b**). Red asterisk indicates the position of Pi binding site. **c,** Distribution of positively-charged residues (shown as sticks) along the Pi translocation pathway, with those positioned in the entrance of EV and IV colored green, and those surrounding Pi binding site colored yellow. **d**, Expanded view of the bound Pi and its interaction with the surrounding residues. Two views are shown. **e**, Time-dependent r.m.s.d. of bound Pi during the 200-ns simulations. For each of two bound Pi in a XPR1 dimer, the r.m.s.d. values of three replicates are overlaid and plotted separately. **f**, Relative Pi export level in cells expressing Pi coordinating residues mutants compared with Pi export level measured in XPR1 WT cells at time point 30min. n=5 independent experiments. One-way ANOVA with Dunnett’s multiple comparisons test **** = p ≤ 0.0001.

In support of the claim about the Pi binding site, an extra density is observed at this site in both structures, likely representing bound Pi molecule (Supplementary Fig. 2). Multiple residues, mostly positively-charged ones, surround the bound Pi and are likely involved in forming interactions with the bound Pi. These residues include Asp 398 and Asn 401 from TM 5, Arg 459 from TM6, Lys 482 and Tyr 483 from TM7, Asp 529 from TM8, Arg570 from TM9, and Arg 603 and Arg 604 from TM10, which are highly conserved among XPR1 homologs (Fig. 3d). Pi in solution are known to exist in different forms according to pH, e.g., dibasic (HPO_4_^2-^) at pH ∼8, or monobasic (H_2_PO_4_^-^) at acidic pH^30^. Since our structures were obtained at ∼pH 7.4, bound Pi is likely in dibasic form. However, as the exact position of the hydrogen group and oxygen groups in Pi could not be defined due to limit in resolution of our XPR1 structures, specific interactions involved in Pi binding could not be depicted. Nevertheless, large number of hydrophilic residues suggest a binding mode involving extensive hydrogen bonding network, similar to the case in the soluble phosphate binding protein (PBP) of bacterial ABC-type high affinity phosphate transport system (pstSCAB complex, whose PBP is pstS), in which up to 13 hydrogen bonds are formed between protein and Pi to achieve high specificity and affinity (Supplementary Fig. 6a)^31^. In addition, positively charged Arg 135 in pstS not only neutralizes the negative charge in Pi, but also forms hydrogen bonds with O2 group of Pi, whereas negatively charged Asp 56 accepts a proton on Pi (Supplementary Fig. 6a)^31^. However, given that much more positively-charged residues are present in Pi binding site of XPR1 when compared with that of pstS (5 in XPR1 vs 1 in pstS), differences in binding mode are expected and await further investigations in the context of Pi bound XPR1 structures at higher resolution. Interestingly, configuration of Pi binding site in XPR1 is more similar to that of Glycerol-3-Phosphate (G3P) Transporter (GlpT), which is an organic phosphate/(Pi) antiporter responsible for G3P uptake^32^. In both XPR1 and GlpT, Pi-coordinating residues are dominated by positively charged residues (Arg 45 and Arg 269 In GlpT) (Supplementary Fig. 6b)^32^. Moreover, based on biochemical data, similar Pi binding environment is likely also present in bacterial high affinity Pi importer PstSCAB, as multiple positively-charged residues appear to be essential for Pi transporting (Arg 170 and Arg 220 in pstA, and Arg 237 in pstC), intertwined with negatively-charged residues (Glu 173 in pstA and Glu 240 in pstC) and other polar residues ^33,34^.

In support of their key roles in Pi binding, these residues are highly conserved among XPR1 homologs (Supplementary Fig. 7). In addition to their role of charge neutralization in Pi binding, these positive-charged residues might also play a role in repulsion of cations and therefore contribute to ion selectivity of XPR1 towards negatively-charged Pi. Besides a proper chemical environment, Pi binding at this site could also be supported by MD simulations, during which modeled Pi remained relatively stable (Fig. 3e). Moreover, mutation of selected residues either abolished or largely impaired the Pi export activity in our assay system, without significantly affecting their cell surface expression pattern, as demonstrated using flow cytometry (Fig. 3f, Supplementary Fig. 5&8).

### Resting conformation of XPR1 and possible mechanism of Pi translocation

In the absence of PP-IPs, Pi export is impaired in human cells despite the presence of XPR1 at the plasma membrane, suggesting that XPR1 is in a non-conducting state. In order to capture XPR1 in an inactive conformation, we generated a triple mutation in the SPX domain (Y22A, E23A and K26A, referred to as XPR1^3m^^ut^ hereafter), which presumably abolished PP-IP binding^13^. After expression and purification in a condition similar to XPR1^wt^, we obtained its cryo-EM structure at a resolution of 2.6 Å (Supplementary Fig. 9).

When comparing the structures of XPR1^3m^^ut^ and XPR1^wt^, the EV of XPR1^wt^ which is wide open, becomes largely collapsed in XPR1^3m^^ut^, mainly due to structural changes in TM9 and the loop between TM9 and TM10 (Fig. 4a-b). Specifically, C-terminus of TM9 swings towards the center of phosphate translocation pathway by ∼45°, around the residue of Arg 570 (Fig. 4c). As a result, TM9, which is bent in XPR1^wt^, become a straight and continuous helix in XPR1^3m^^ut^ (Fig. 4c). Despite dramatic structural changes in TM9, to our surprise, structure of most other part of XPR1^3m^^ut^ remain unchanged when compared with that of XPR1^wt^, including the IV. The only exception is the extracellular half of TM7, TM 8 and the loop between them, which are slightly pushed outward away from the Pi translocation pathway in XPR1^3m^^ut^ (Fig. 4c), likely an accompanying event of movement of the C-terminal half of TM9.

**Figure 4.**
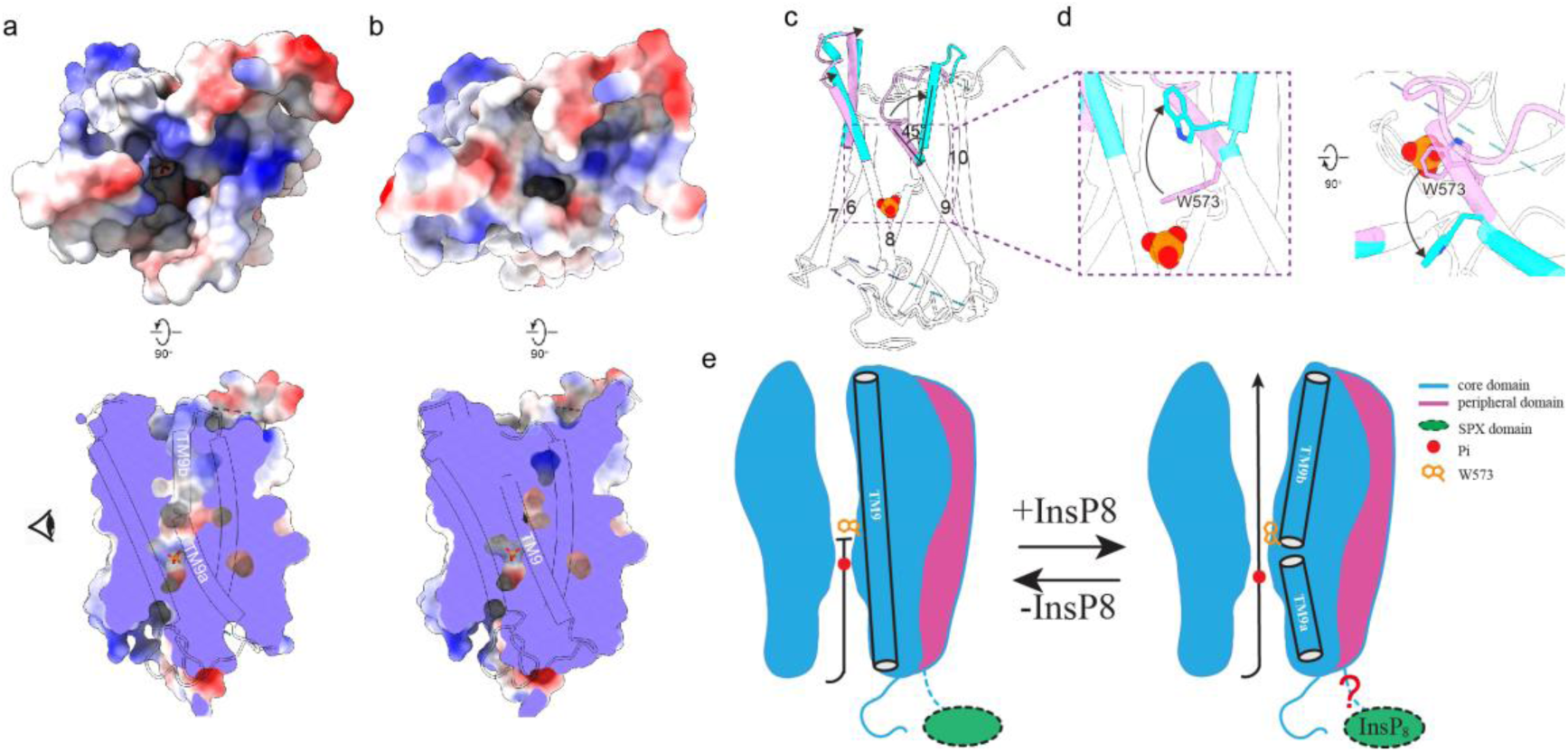
Structure of XPR1^3m^^ut^, its comparison with XPR1^wt^, and proposed mechanism. **a-b**, Comparison of surface electrostatic potential of XPR1^wt^ (**a**) and XPR1^3m^^ut^ (**b**), highlighting the changes in the EV. Two views are shown for each structure. **c,** Superimposition of structures of XPR1^wt^ and XPR1^3m^^ut^, highlighting the conformational changes occurred in extracellular half of TM9 (cyan for XPR1^wt^ and purple for XPR1^3m^^ut^), viewed from the angle indicated in **a** (eye symbol). **d**, Expanded view of the region boxed in **c** in two views, highlighting the movement of sidechain of Trp 573. **e,** Schematics of proposed working model. Only one protomer is shown, with each domain represented by different shapes and colored differently in the same way as in Fig. 2a.

Zooming in into the region around the Pi binding site, we found that in XPR1^3m^^ut^, Trp 573 in TM9 sits right above the bound Pi, likely preventing it from leaving the binding pocket and entering into the EV (Fig. 4d). By contrast, in XPR1^wt^, sidechain of Trp 573 is dramatically rearranged, along with other part of TM9b that undergoes large conformational changes (Fig. 4d). Specifically, side chain of Trp 573 in XPR1^wt^ is lifted upward and swings towards the entrance of EV, thereby unleashes the caged Pi and allows them to enter into the EV, and eventually out of cell (Fig. 4d). Therefore, we propose that Trp 573 functions as an extracellular gate and controls the release of bound Pi in the Pi binding pocket (Fig. 4e). However, since the SPX domain is not resolved in both structures of XPR1, how binding of PP-IPs in the SPX domain causes conformational changes in Pi translocation pathway remains to be addressed.

## Discussion

Despite the critical importance of phosphate in health and disease, our understanding in the molecular basis of Pi export is poor, owing to the lack of a three-dimendional structure of XPR1 required for structure-function investigations. In this study, we determined cryo-EM structures of human XPR1, a recently identified Pi exporter, in two distinct conformations (Fig. 1&4). We found that XPR1 forms a homodimer, with the core domain of each protomer being distantly related to ion-translocating rhodopsin/channelrhodopsin (Fig. 2). An ion translocating pathway is observed in the core domain, with a Pi molecule bound in the narrowest point in middle of the membrane. These structural observations, along with in vitro Pi export assays and computational analysis, confirm the functional role of XPR1 as Pi exporter (Fig. 3). By comparing its two structures, which are likely in closed and conductive state respectively, a possible working model of Pi export by XPR1 has been proposed (Fig. 4). Although a lot of questions remain to be addressed, our study provides a framework for future in-depth mechanistic studies of this pharmaceutically relevant protein.

We noticed that, a highly conserved stretch of residues (Asn 618 - Asp 628) located at the C-terminus of TM10 deviates away from other part of TM10 and forms an ordered and expanded loop structure, which appears to partially block the intracellular entrance of phosphate translocating pathway in XPR1 (Supplementary Fig. 7, 10, 11). This loop is highly conserved among XPR1 from different animal species. If this loop is manually removed from XPR1 structure, the IV becomes much larger when compared with that of intact XPR1 structure (Supplementary Fig. 10b-c). Surprisingly, among the reported PFBC variants, at least three are located in or around this region, including N619D, R624H and I629S^17,18^, suggesting that this region may play an important role in regulating Pi export. Whether and how this peptide participates in Pi export await further investigations.

In addition to Pi binding proteins mentioned above, structural information is available for a few additional Pi transporters, including those of a fungal (*Piriformospora indica*) high-affinity phosphate transporter PiPT^35^ and of the sodium-dependent phosphate transporter from *Thermotoga maritima* (TmPiT)^36^, to our knowledge. However, chemical environment for Pi binding of these proteins is dramatically different from that of XPR1. Indeed, positively-charged residues are barely involved in Pi binding in PiPT and TmPiT (Supplementary Fig. 6c-d). Instead, negatively-charged as well as other polar residues are found in the Pi binding pocket and directly interact with Pi in PiPT and TmPiT by forming hydrogen bonds with bound Pi (Supplementary Fig. 6c-d). While seemingly unfavorable chemistry-wise, positively-charged counter-ions in PiPT and TmPiT (Proton for PiPT and Na^+^ for TmPiT) ^35,36^ may play the same roles as positively charged residues in proteins that only bind Pi in their substrate binding pocket, such as XPR1, GlpT, and pstSCAB.

Since the SPX domain in both states is not resolved, and more PFBC related mutations reported so far are located in this domain, we went ahead and predicted the full-length model of XPR1, on which the disease-related mutations can be mapped. The XPR1 AlphaFold model of XPR1 is overall quite accurate (a RMSD of 1.08 Å when TMD of dimeric cryo-EM structure and AlphaFold model are superimposed) (Supplementary Fig. 12a-b), except the peptide at the C-terminus of TM10, which was predicted to form a continuous helix instead of an extended loop observed in our cryo-EM structures (Supplementary Fig. 12a). When the disease-related mutations were mapped onto the AlphaFold model of XPR1, it is obvious that SPX domain is the top hotspot, followed by C-terminus of TM10 and ion-translocation pathway (Supplementary Fig. 13), which is consistent with their degree of conservation (higher conservation in intracellular side than extracellular side, see Supplementary Fig. 11).

In summary, our study not only confirms the identity of XPR1 as phosphate exporter and paves the way for further in-depth mechanistic studies of XPR1 and its homologs, but also lays the foundation for future structure based drug design targeting XPR1.

## Data availability

Structure coordinates and cryo-EM density maps have been deposited at the Protein Data Bank and Electron Microscopy Data Bank under accession numbers xxxx and EMD-xxxxx for XPR1^wt^; xxxx and EMD-xxxxx for XPR1^3m^^ut^. Other structure coordinate analyzed in this paper are indicated in the text.

## Author Contributions

Q.C., B.L., and J.B. conceived and supervised the project. Q.H. prepared the samples; R.Z., B.L., and Q.C. performed data acquisition, image processing and structure determination; S.T., V.C., and J.B. performed Pi export assay and analyzed the data; Q.C. performed MD simulations; all authors participated in research design, data analysis, and manuscript preparation.

## Acknowledgements

We thank the staff members of the Center of cryo-EM of Fudan University for technical support and assistance. This work was supported in part by grants from The National Science and Technology Innovation 2030 Major Projects of China (STI2030-Major Projects-2022ZD0207800 to B.L.), the National Natural Science Foundation of China (32071202 and 32271012 to Q.C.; 32371261 to B.L.), Tianjin Fund for Distinguished Young Scholars (20JCJQJC00080 to Q.C.), Yunnan Fund for Distinguished Young Scholars (202401AV070004 to Q.C.), the Xingdian Scholar Fund of Yunnan to Q.C., and the Postgraduate Research and Innovation Foundation of Yunnan University (KC-23236412).

## Methods

### Protein expression and purification

CDS of wild type (XPR1^wt^) and triple mutation (XPR1^3mut^) of Human XPR1 (NCBI refseq XM_013083282.1) fused with a C-terminal strep tag was cloned into a pEZT vector^37^ and heterologously expressed using the BacMam system. The baculovirus was generated in Sf9 cells grown in SIM SF Expression Medium (Sino Biological Inc) following standard protocols using transfection reagent X-tremeGENE™(Roche) and used to infect HEK293F cells grown in SMM 293-TI complete medium (Sino Biological Inc) at a ratio of 1:40 (virus:HEK293F, v:v). 10 mM sodium butyrate was supplemented to boost protein expression. Subsequent purification steps were carried out at 4°C.

48 hours after infection, cells were harvested by centrifugation at 4,000g. The cell pellet was resuspended in buffer A (50 mM Tris pH 8.0, 200 mM NaCl), supplemented with 1mM PMSF and disrupted by sonication on ice. For XPR1^wt^ but not XPR1^3mut^, we isolated the cell membrane first. In brief, the lysate was centrifuged at 40,000 RPM for 1 h to pellet membranes, which were then Dounce homogenized in buffer A supplemented with 1 mM PMSF. Protein was extracted from isolated membrane (XPR1^wt^) or whole cell lysate (XPR1^3mut^) with 1% (w:v) DDM by gentle agitation for 2 hours. Insoluble material was removed by centrifugation at 20,000 RPM for 40 minutes and the supernatant was incubated with Strep-Tactin®XT resin (IBA Lifesciences) for 2 hours with gentle agitation. The resin was then collected in a disposable gravity column (Bio-Rad), washed with buffer A supplemented with 0.06% (w:v) GDN, and finally eluted with the same buffer containing biotin. The eluant was concentrated, and further purified by size exclusion chromatography on a Superose 6 10/300 GL column (GE Heathcare) pre-equilibrated with PBS buffer supplemented with 0.06% (w:v) GDN. The peak was collected and concentrated to ∼4 mg/ml for Cryo-electron microscopy analysis.

### Grid preparation

The XPR1^wt^ and XPR^3mut^ samples were centrifuged at 21,000 x g for 10 min at 4°C before grids preparation. A 3.5 µL sample was applied to holey carbon, 300 mesh R1.2/1.3 gold grids (Quantifoil, Großlöbichau, Germany) that were freshly glow discharged in H2-O2 mixture for 30-60 seconds. Sample was incubated for 5 seconds at 4°C and 100% humidity prior to blotting with Whatman #1 filter paper for 3-3.5 seconds at blot force 1 and plunge-freezed in liquid ethane cooled by liquid nitrogen using a FEI Mark IV Vitrobot (FEI / Thermo Scientific, USA).

### Cryo-EM data acquisition and data processing

All datasets collected on Titan Krios G4 cryo-electron microscope operated at 300 kV, equipped with a Falcon G4i direct electron detector with a Selectris X imaging filter (Thermo Fisher Scientific) operated with a 20-eV slit. Movie stacks were automatically acquired using the EPU software (Thermo Fisher Scientific) in super-resolution mode with pixel size of 0.466 Å with a defocus range of −0.8 to −1.6 μm. The total dose per EER (electron event representation) movie was 47.66 e^-^/Å and 49.97 e^-^/Å for XPR1^wt^ and XPR1^3mut^, respectively.

For XPR1^wt^ dataset, 3,014 micrographs were collected. Each EER movie with 1,080 frames were fractionated into 40 subgroups, and beam-induced drift was corrected using MotionCor2^38^ in Relion (v4.0.0)^39^. The data was binned to 0.932 Å/pixel. After the autopicking and rounds of 2D classification, 608,818 particles were imported into cryoSPARC (v4.0.2)^40^ for the further 2D classification. Good classes were used to generate an ab initio model with 2 classes and 0 similarity with or without symmetry for couple times until good model generated. Particles belonging to a class with well-defined features were further refined using Heterogeneous refinement and Non-uniform (NU) refinement with C2 symmetry, which finally yielded a map at 2.9 Å for model building.

For XPR1^3m^^ut^, total of 4,300 EER movie stacks were collected and processed in a similar way as XPR1^wt^ dataset. In order to gain more side view information, we selected the classes with good side view to train the micrographs with Topaz^41^ for particle picking, and then merged with previous particles for the further processing. Total of 247,144 particles were futher refined using Non-uniform (NU) refinement with C2 symmetry and yielded final maps at 2.6 Å.

### Model building, refinement, and validation

Model building was manually conducted in Coot^42^, guided by Alphafold predicted model (AF-Q9UBH6-F1). Accurate residue registration was achieved based mainly on the clearly defined densities for bulky residues (Phe, Trp, Tyr, and Arg). Models were refined against cryo-EM maps using real-space refinement in PHENIX ^43^, with secondary structure and non-crystallography symmetry restraints applied. For modeling and refinement of the POPC molecule, its SMILE strings was inputted into eLBOW ^44^ implemented in PHENIX to generate a CIF file. The statistic of the models’ geometries was generated using MolProbity ^45^. Ion translocating pathway was calculated using the CAVER program ^46^. All the figures were prepared in PyMol^47^ or ChimeraX^48^.

### Cell surface monitoring of XPR1

HCT116 (human colorectal cancer cell line, ATCC CCL-247) were maintained in DMEM (ThermoFisher scientific) supplemented with 10% fetal bovine serum (FBS, Sigma), 1% antibiotics (penicillin-streptomycin, Sigma) and non-essential amino acids, in a 5% CO_2_ incubator at 37°C. Cells were detached from plates using 0.05% of trypsin and washed with cold PBS supplemented with 2% FBS and 1 mM EDTA (PBAE). 10^5^ cells per well in 96-well plate (U bottom) were then incubated for 30’ at 4°C in PBAE containing either the previously described XPR1 immunoadhesin ligand XRBD (xenotropic Env receptor-binding domain fused to the mouse IgG1 Fc)^12^ or a mock control. Cells were then washed twice with cold PBAE and labeled for 20 min on ice with Alexa 647-conjugated anti-mouse IgG1 antibodies (1:250, Invitrogen) in PBAE. Cells were then washed in PBAE and analyzed by flow cytometry on a Novocyte (Acea, Biosciences, Inc). Data analysis was performed using FlowJo software.

### Generation of XPR1 KO cells

HCT116 cells were transiently transfected (Jet Prime transfection reagent, Polyplus) with the PX458 plasmid (addgene #48138) ^49^ containing the Cas9 gene, the single guided RNA targeting the first exon of the *XPR1* gene (5’-GCCCTCACTACACGCGCCTC-3’) and the green fluorescent protein *GFP* gene. Two days after transfection, GFP-positive cells were purified by flow cytometry and grown for 1 week in culture medium. Cells lacking cell surface XPR1 were sorted by flow cytometry using the XRBD ligand and several individual XPR1 KO cell clones were isolated, tested for the absence of XPR1 and used for further analyses.

### Phosphate export assay

Confluent cells in a 12-well plate were incubated with fresh DMEM 10% FBS for 1 hour at 37°C and placed on ice for 30’ before washing three times with cold phosphate-free DMEM. Cells were the incubated for the indicated periods of time in phosphate-free DMEM supplemented with 10% dialyzed FBS. Supernatant were then harvested, centrifuged 3’ at 5000 rpm, and presence of phosphate was detected using the malachite green phosphate assay kit (MAK307, Sigma). Cells were lysed in 0.5% SDS and concentrations of proteins were determined using the BCA protein assay kit (Pierce) for further normalization.

### MD simulations

MD simulation systems for the XPR1 structures were built using the CHARMM-GUI server ^50^, with each structure being embedded in a palmitoyl-oleyl-phosphatidylcholine (POPC) bilayer hydrated with water molecules (using TIP3P model). 700 mM NaCl was added to each system. All simulations were carried out with the Gromacs package 2021.4 ^51^ using the CHARMM36m force field ^52^. Following 50,000 steps of energy minimization with the steepest descent algorithm, a 1,125 ps of equilibration at a constant temperature of 300 K was performed, during which positional restraints were gradually reduced, and Berendsen thermostat and barostat were applied. During production runs, temperature and pressure were controlled with a Nose-Hoover thermostat and a Parrinello-Rahman barostat.

## Supplementary figures and legend

**Supplementary Fig. 1.**
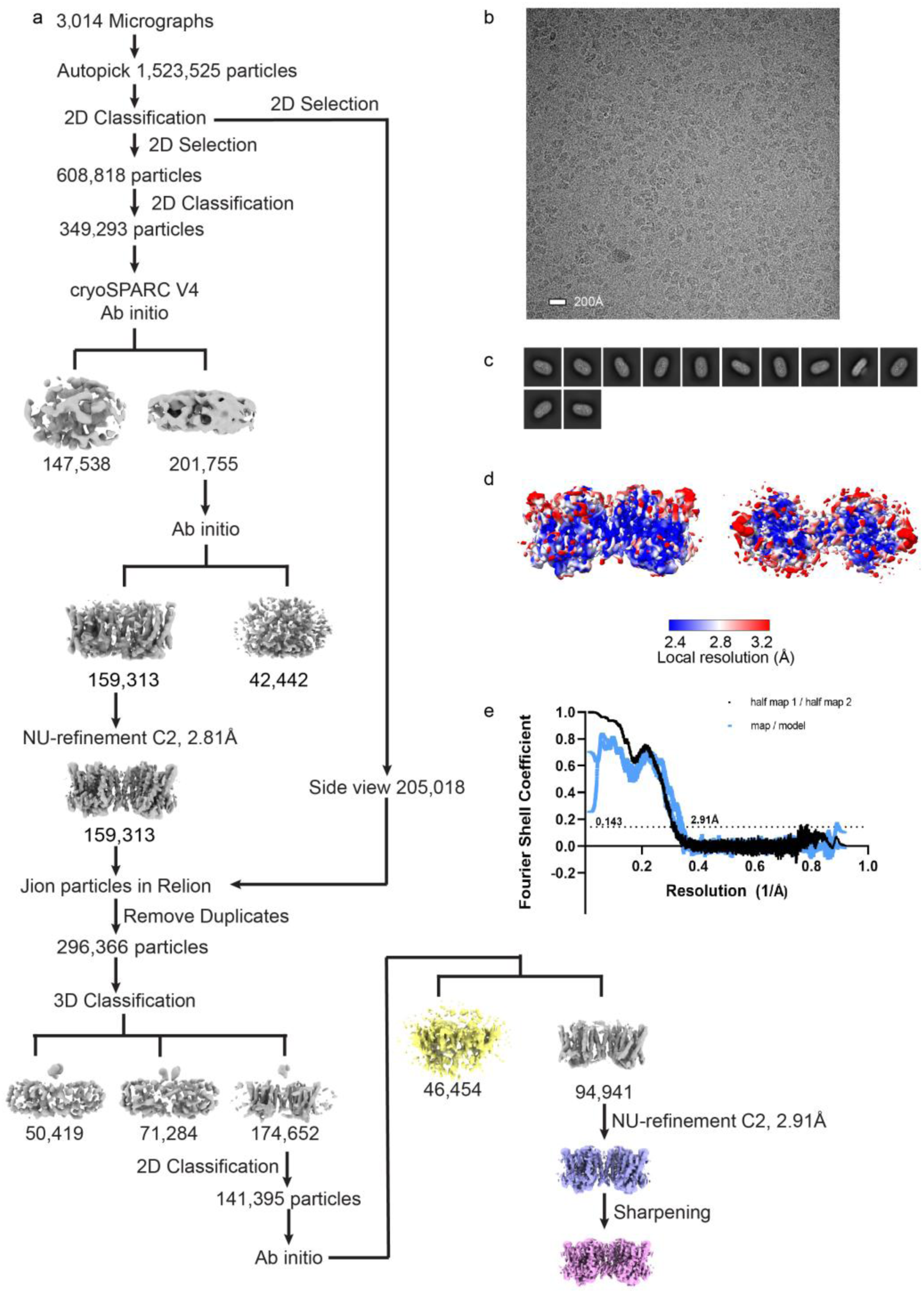
Cryo-EM analysis of XPR1^wt^. **a,** Workflow of cryo-EM image processing and reconstruction. **b,** A representative cryo-EM micrograph. Scale bar, 20 nm. **c,** 2D class average images. **d,** Local resolution distribution. **e,** The GSFSC curve for the reconstruction.

**Supplementary Fig. 2.**
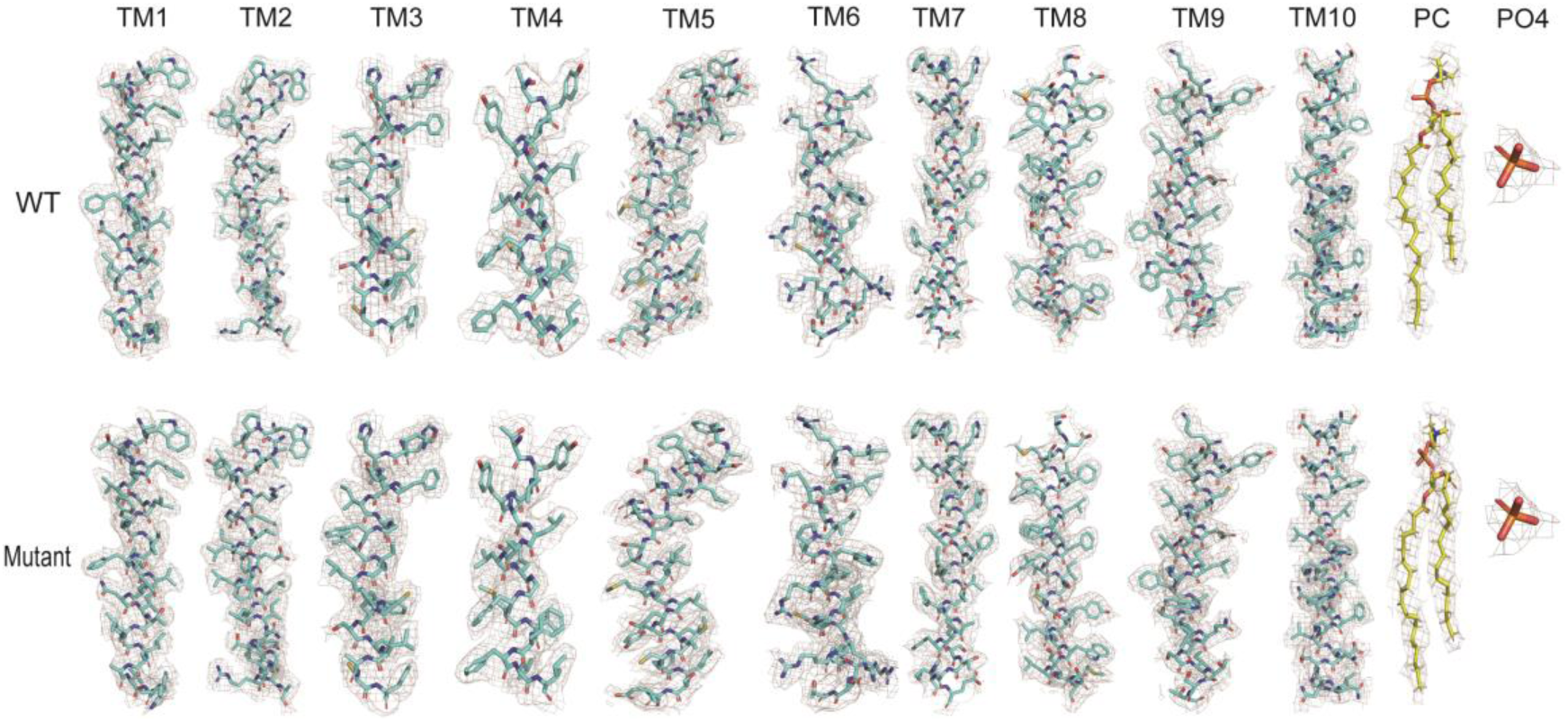
Sample cryo-EM density maps for XPR1^wt^ and XPR1^3m^^ut^. The maps are low-pass filtered to 3Å and sharpened with a temperature factor of −100 Å^2^.

**Supplementary Fig. 3.**
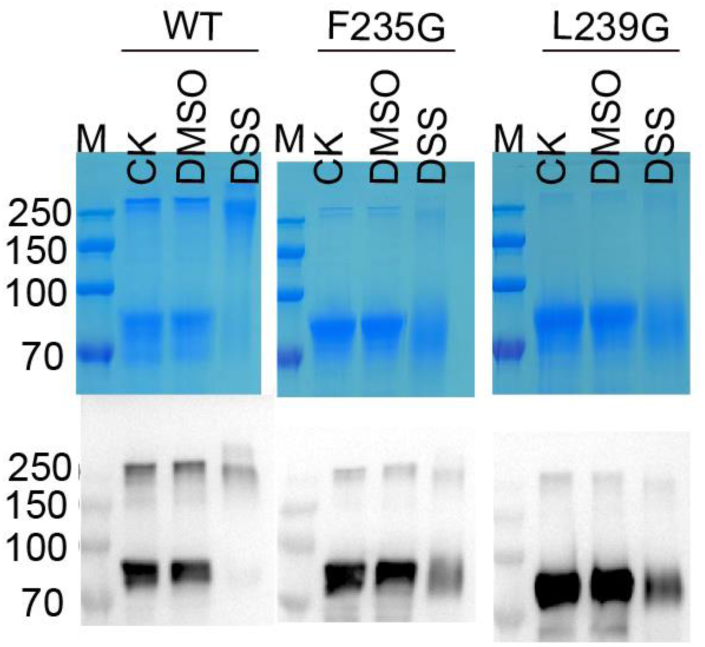
SDS-PAGE (upper) and WB (lower) of crosslinking experiment of wild type XPR1 and its dimer interface mutants. 1 mM DSS was used for crosslinking for 60 min. Note that while monomer of wild type XPR1 can be crosslinked into dimer, it is not the case for mutants.

**Supplementary Fig. 4.**
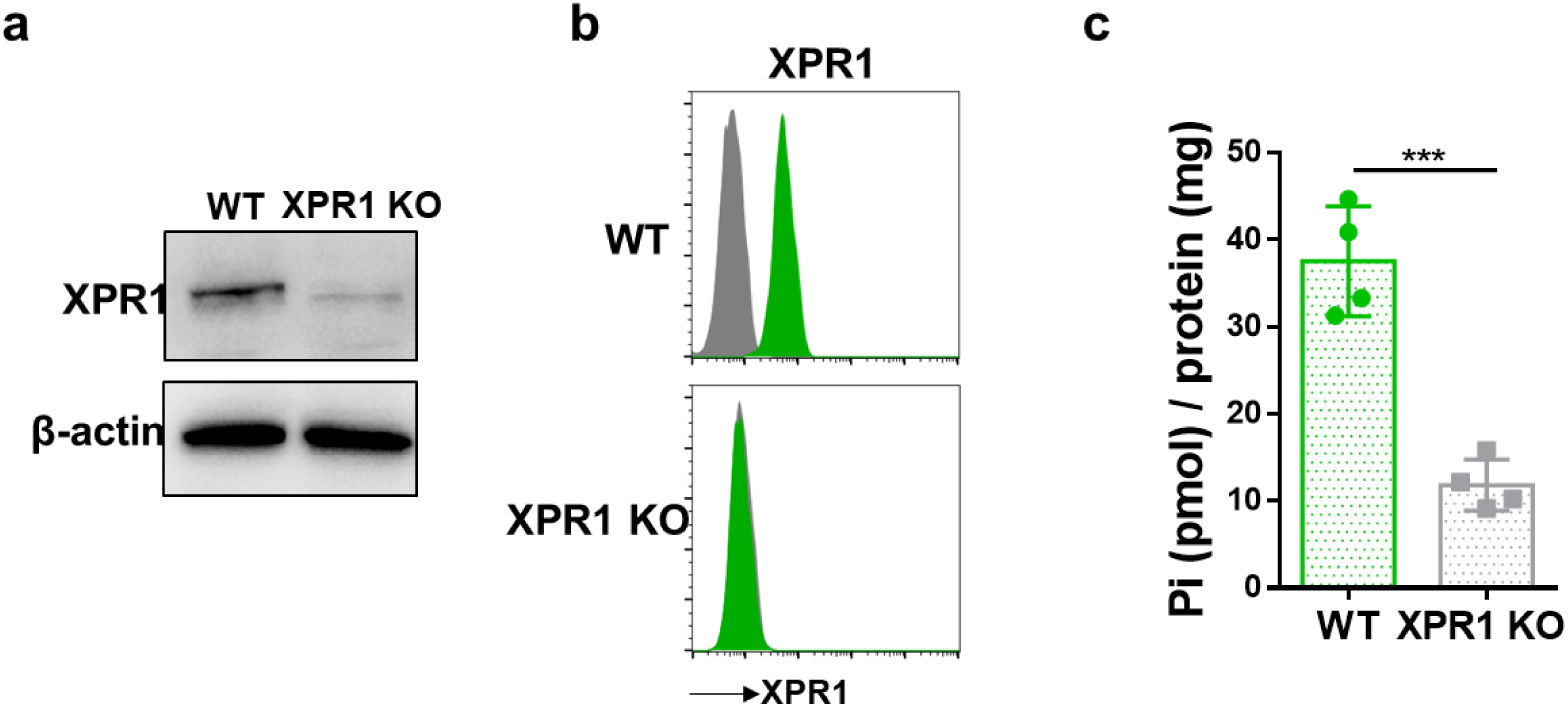
Generation and characterization of XPR1 KO cells. **a.** of XPR1 by immunoblot in cell lysates of parental HCT116 cells (WT) or invalidated for XPR1 (XPR1 KO) generated by CRISPR/cas9 technology. β-actin loading control is shown. **b.** Cells from (a) were evaluated for X-RBD binding by flow cytometry. Curve of mean fluorescence intensity (green) of the XPR1 ligand is compared with non-specific mock staining (grey) of a representative experiment (n=3). **c.** Level of inorganic phosphate (Pi) export from cells from (a) was quantified by malachite green phosphate assay and normalized to the total amount of protein at time point 30min. Data are means ± SD from n=4 experiment. Student unpaired t test *** =p <0.001.

**Supplementary Fig. 5.**
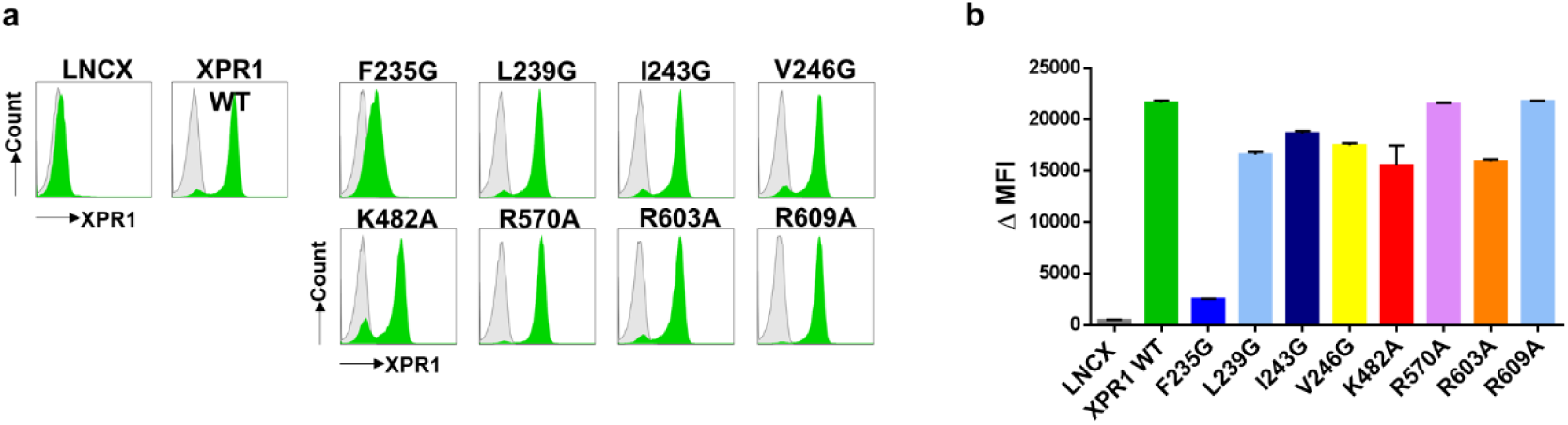
Cell surface expression of XPR1 mutants. **a.** HCT116 XPR1 KO cells and mutants were evaluated for X-RBD binding by flow cytometry. Curve of mean fluorescence intensity (green) of the XPR1 ligand is compared with non-specific mock staining (grey) of a representative experiment (n=3). **b.** Delta mean fluorescence intensity (ΔMFI) bars represents the mean of n=4 independent flow cytometry experiments.

**Supplementary Fig. 6.**
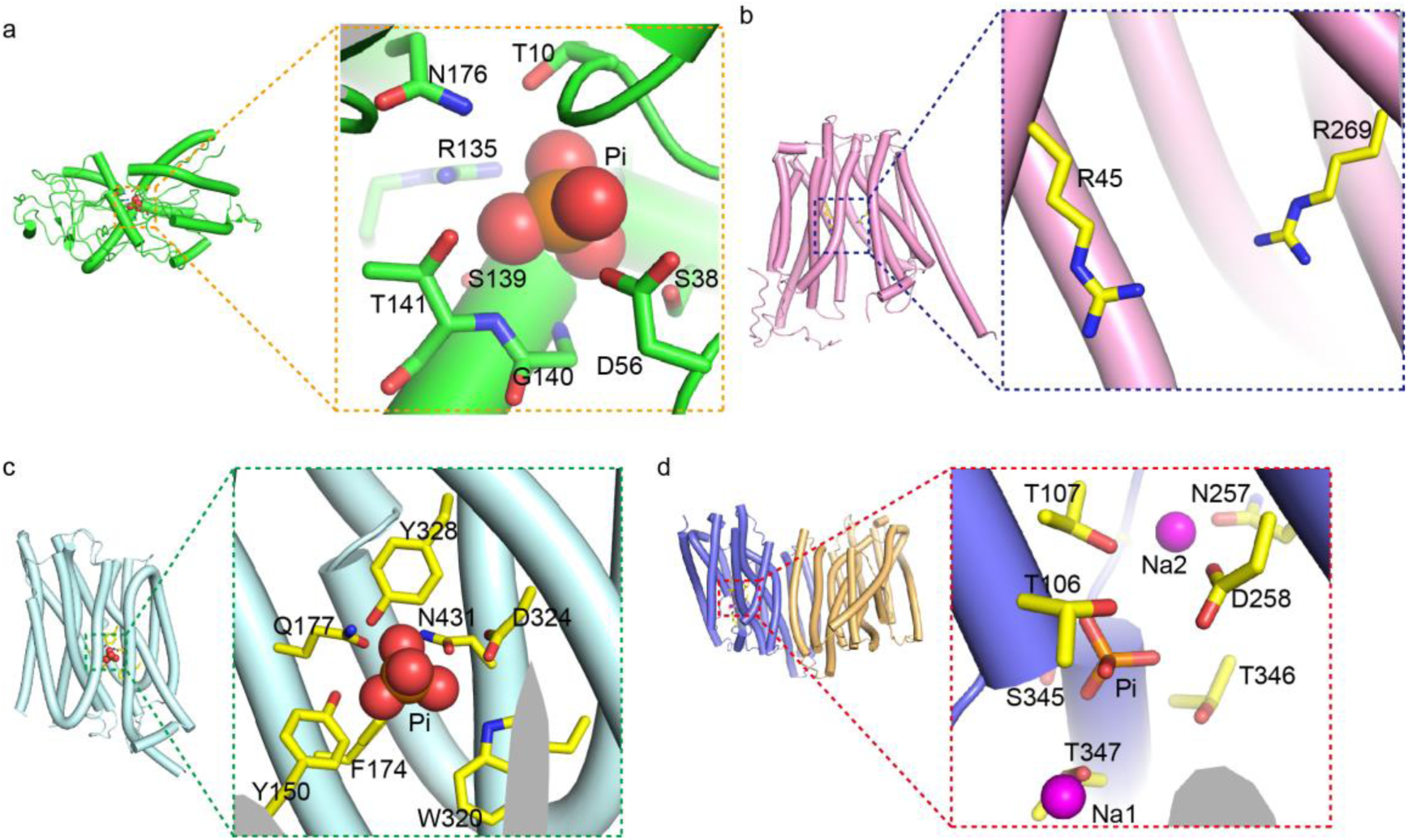
Pi binding sites in pstS (a, pdb #1bpb), GlpT (b, pdb# 1pw4), PiPT (c, pdb# psp5) and TmPiT (d, pdb# 6l85). In each panel, the overview is shown on the left and the expanded view of Pi binding pocket is shown on the right.

**Supplementary Fig. 7.**
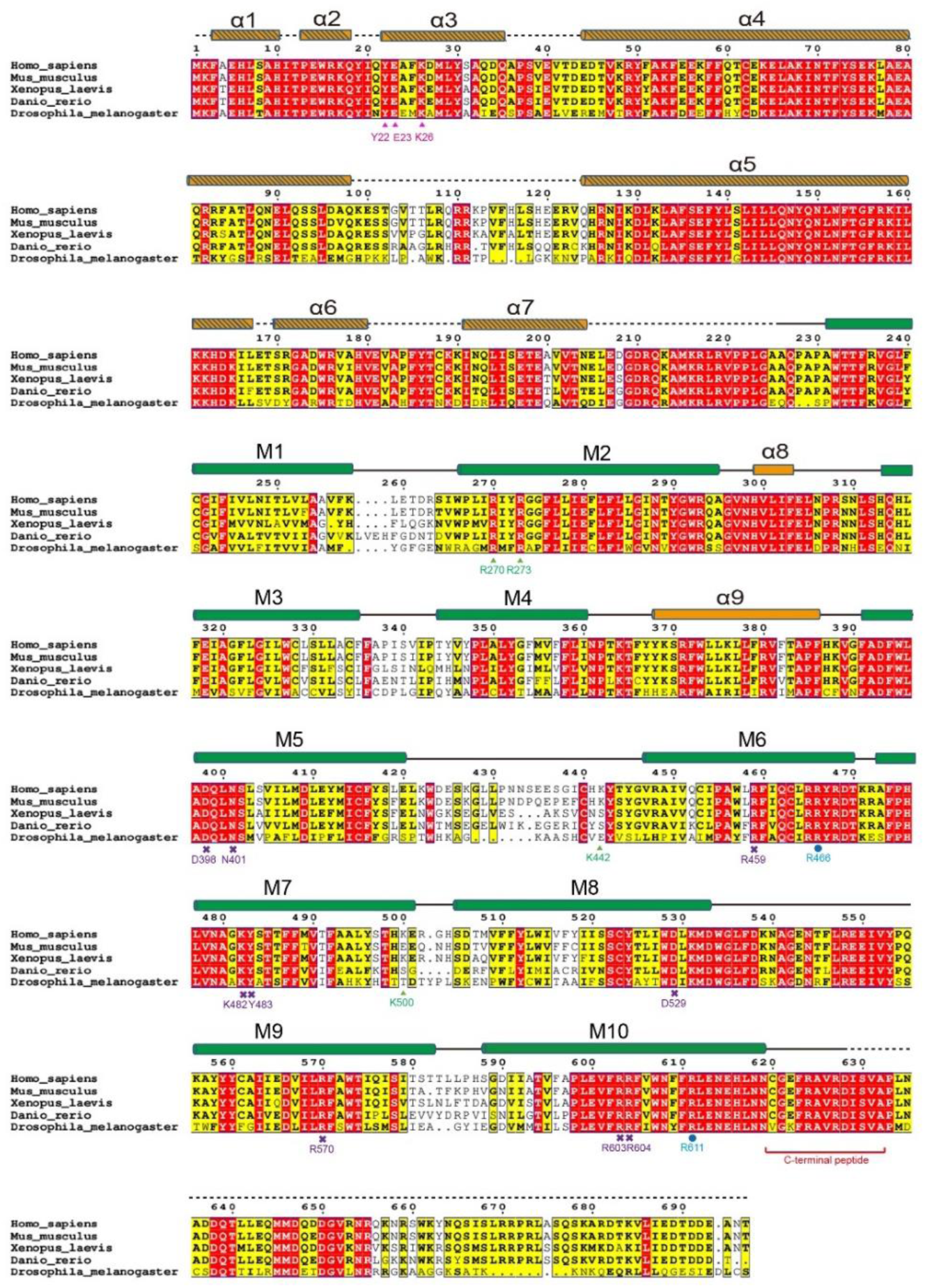
Sequence alignment of XPR1 from representative species. Sequence alignment of human (accession number NP_004727.2), mouse (accession number NP_035403.1), frog (accession number NP_001086930.1), zebrafish (accession number NP_001119862.1) and fruit fly (accession number AAF50730.1) XPR1. Residues involved formation of the Pi binding site are indicated by purple crosses, whereas positively-charged residues in EV and IV are indicated by green triangles and blue circles respectively. α helices of the SPX domain are not visible in our structures and are therefore shown as shaded cylinders. Residues in the possible InsP_8_ binding pocket that were subjected to mutagenesis to generate XPR1^3m^^ut^ are indicated by pink triangles.

**Supplementary Fig. 8.**
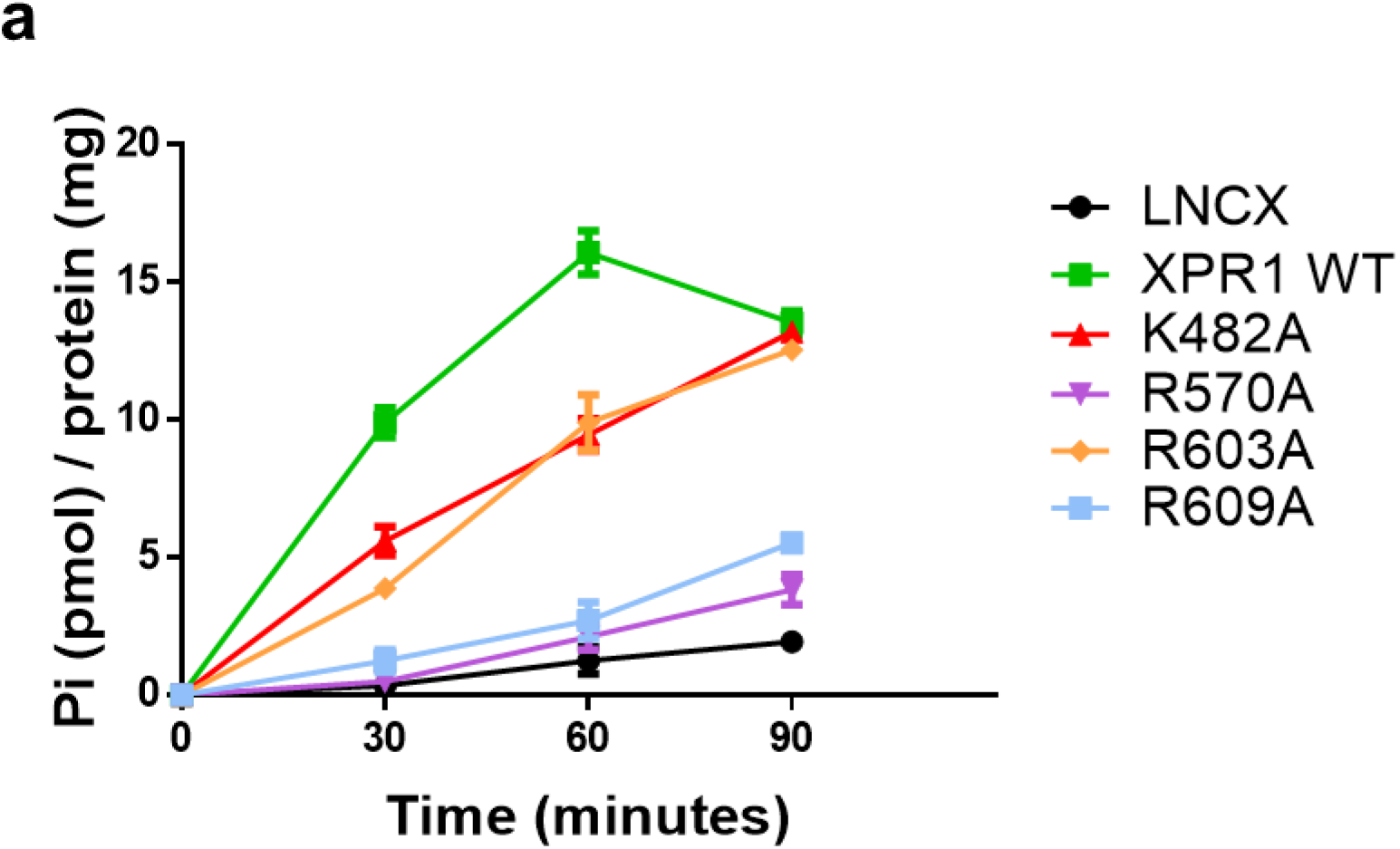
Kinetics of Pi export from HCT116 XPR1 KO introduced with XPR1 WT or mutants on Pi coordinating residues (K482A, R570A, R603A, R609A). Exported Pi was quantified by malachite green phosphate assay and normalized to the total amount of proteins. Data are means ± SD from a representative experiment (n=5).

**Supplementary Fig. 9.**
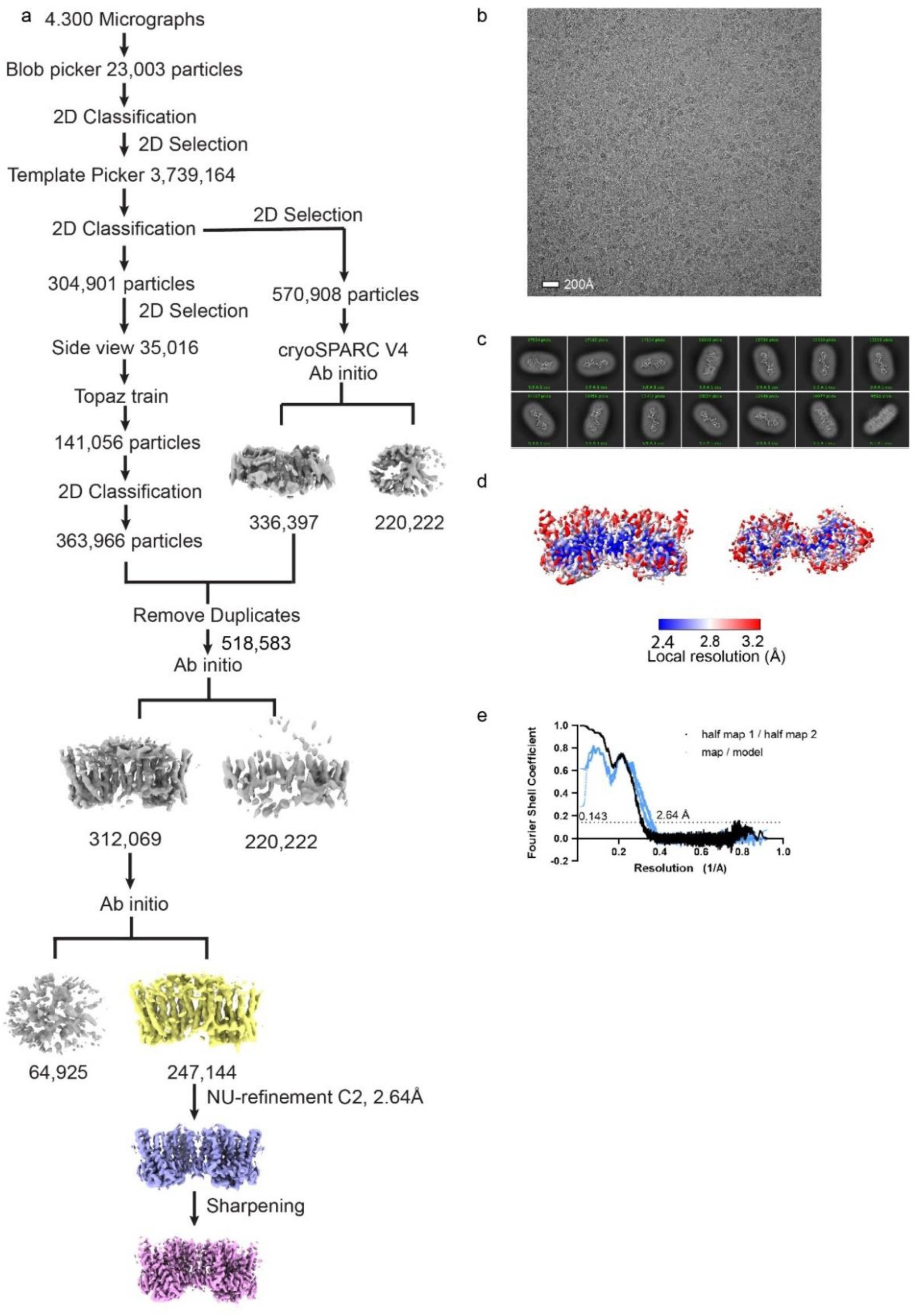
Cryo-EM analysis of XPR1^3m^^ut^. **a,** Workflow of cryo-EM image processing and reconstruction. **b,** A representative cryo-EM micrograph. Scale bar, 20 nm. **c,** 2D class average images. **d,** Local resolution distribution. **e,** The GSFSC curve for the reconstruction.

**Supplementary Fig. 10.**
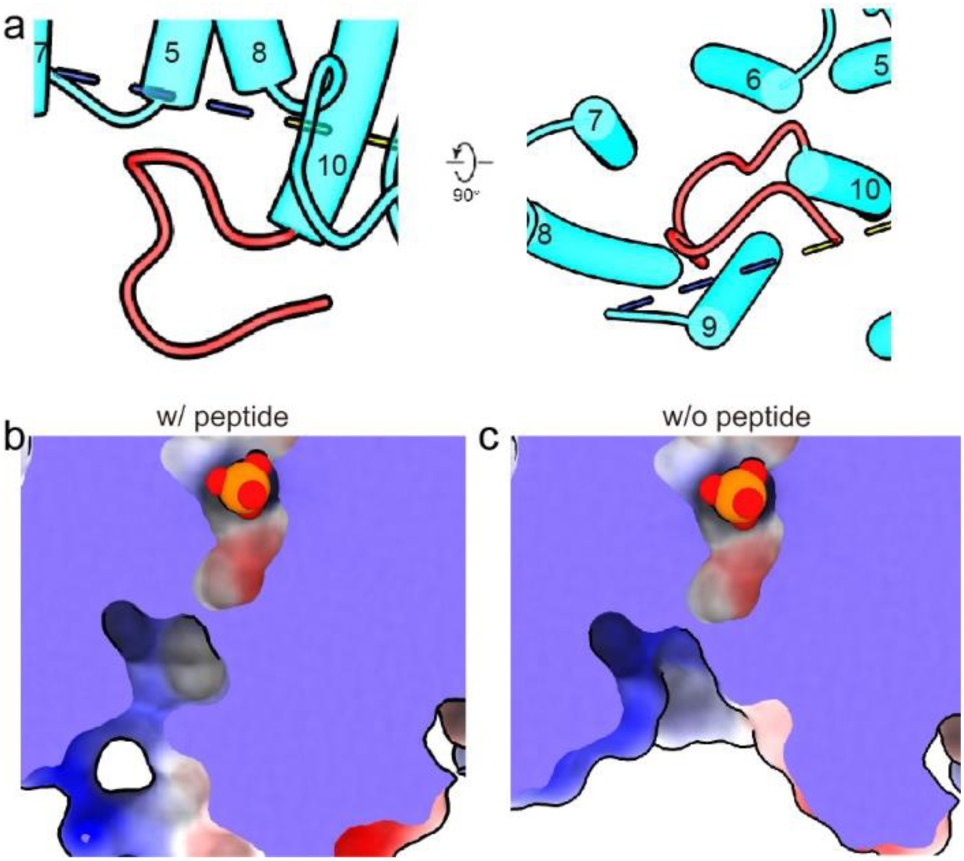
A C-terminal peptide shapes IV of Pi translocation pathway in XPR1. **a**, Location and structure of the blocking peptide in front and bottom views. **b-c,** Difference in the IV between full-length XPR1^wt^ structures (**b**) and XPR1^wt^ model with the C-terminal peptide manually removed (**c**).

**Supplementary Fig. 11.**
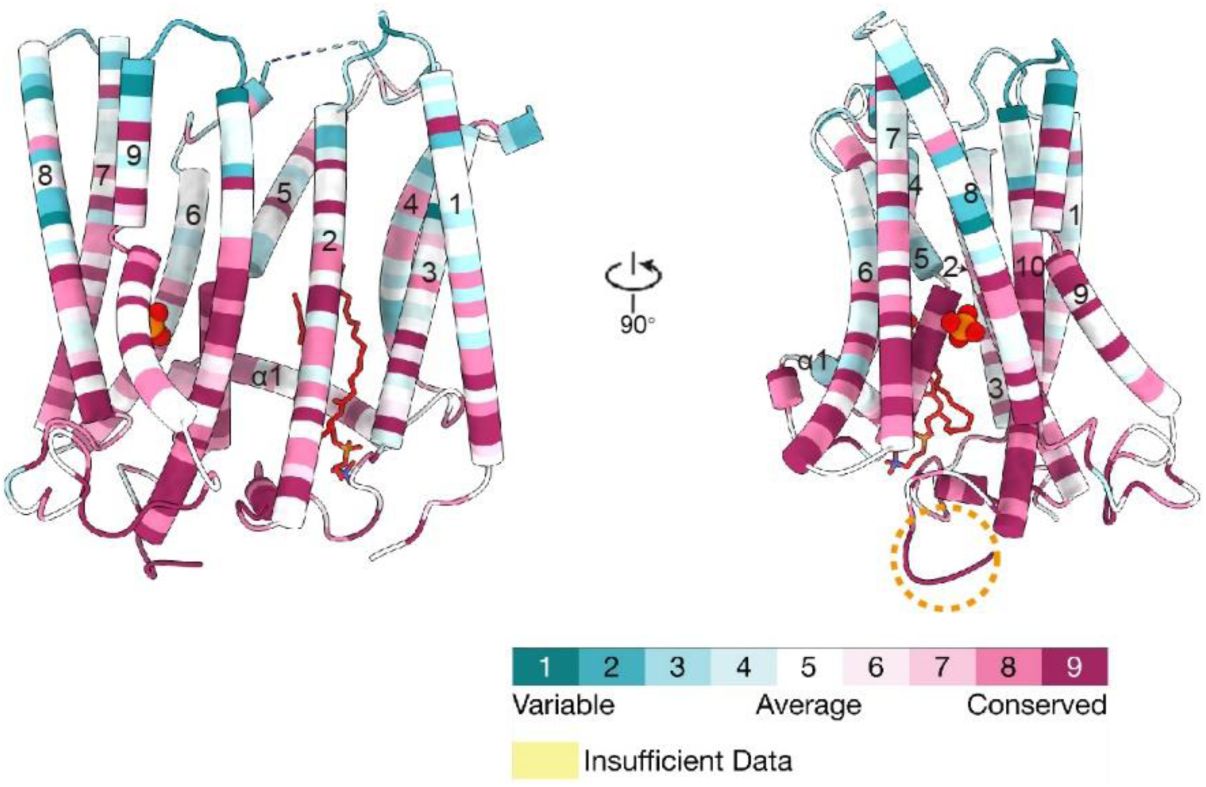
Cartoon representation of XPR1 protomer structure in two views, with each residue colored by its degree of conservation according to MSA shown in Supplementary Fig. 6. Color code is indicated at the bottom. Dashed orange circle highlights the loop at the C-terminus of TM10.

**Supplementary Fig. 12.**
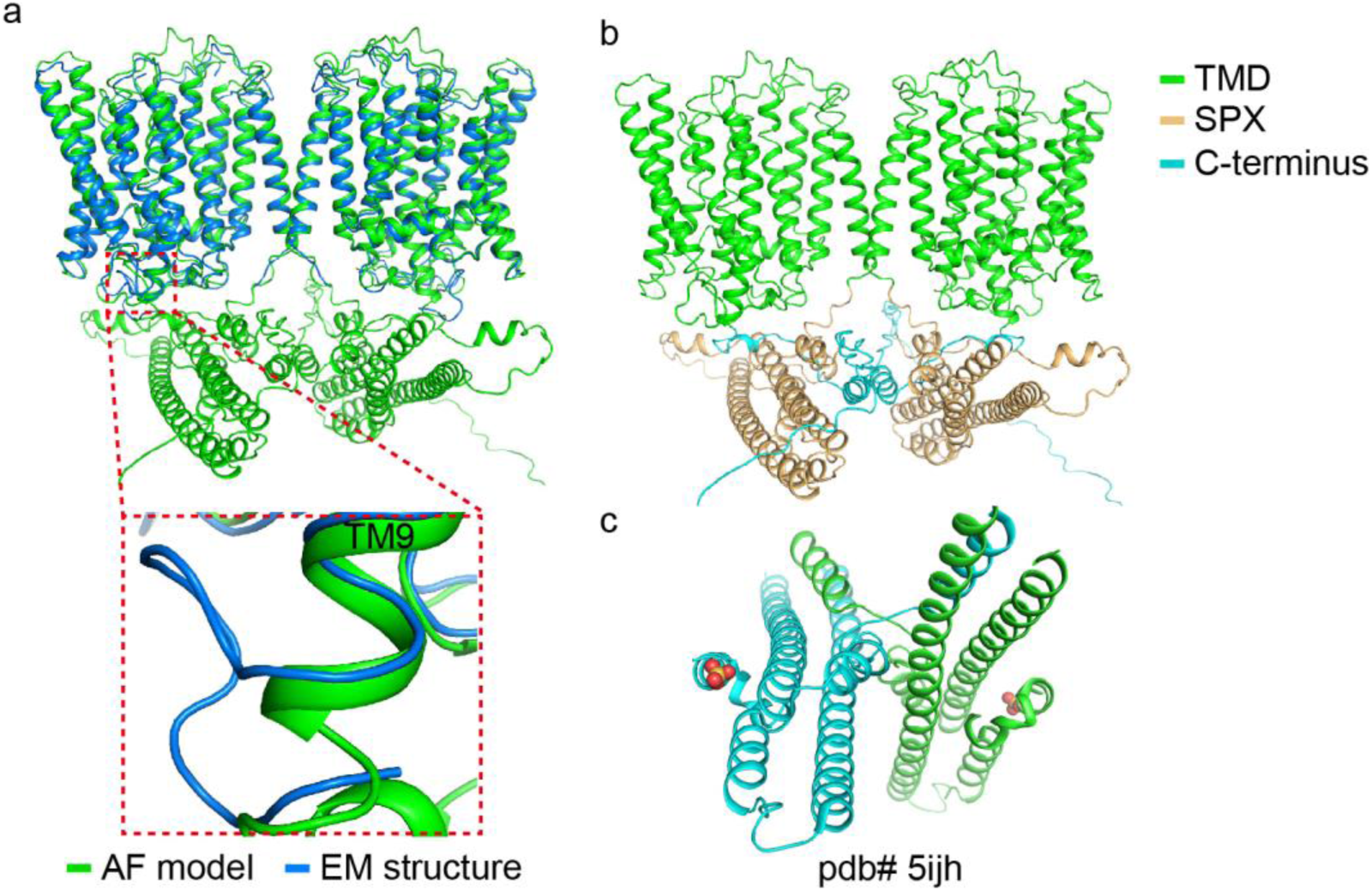
AlphaFold model of XPR1 and its comparison with experimental structures. **a**, Superimposition of AlphaFold model and cryo-em structure of XPR1, with the C-terminus of TM10 highlighted in an expanded view (dashed box). **b,** AlphaFold model of XPR1, with different domains colored differently (see color code). **c,** X-ray crystal structure of human XPR1 SPX domain alone, with the bound sulfate molecules shown as spheres.

**Supplementary Fig. 13.**
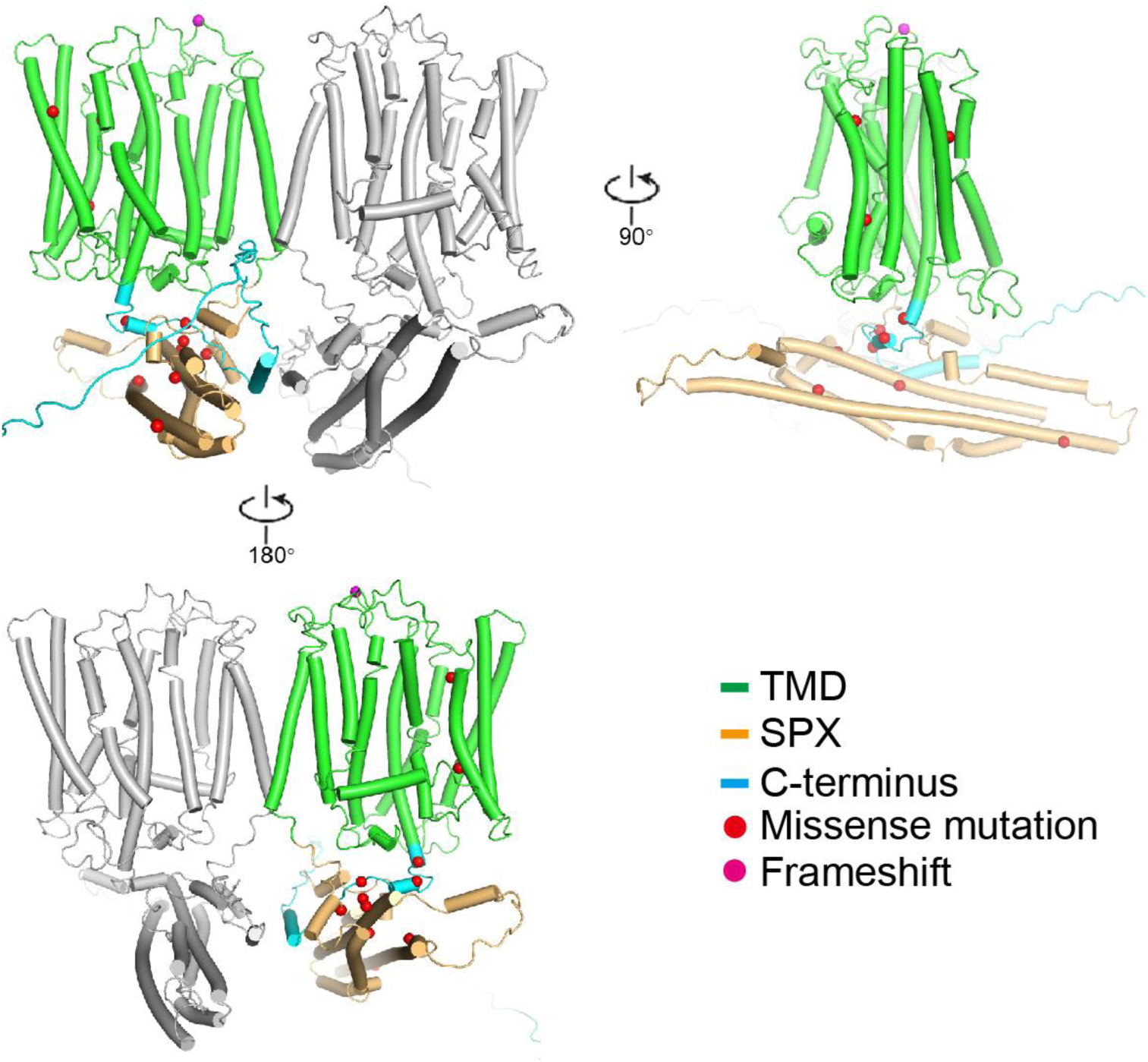
Mapping of PFBC-causing mutations on XPR1 structure model. Dimer is shown, with mutations mapped to one of the monomers. Domains are colored differently and type of mutations are shown as spheres in different color (see color code).

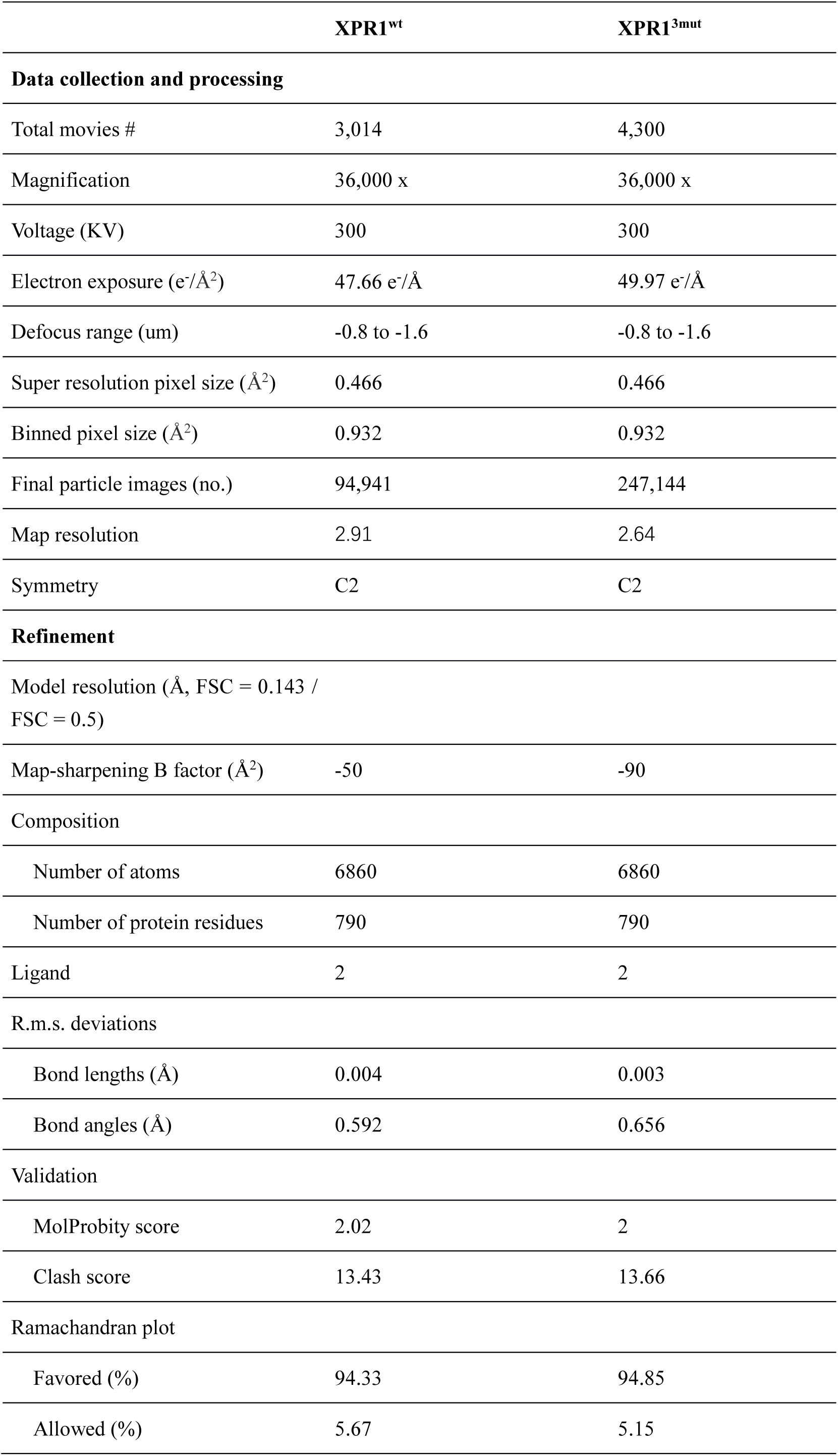
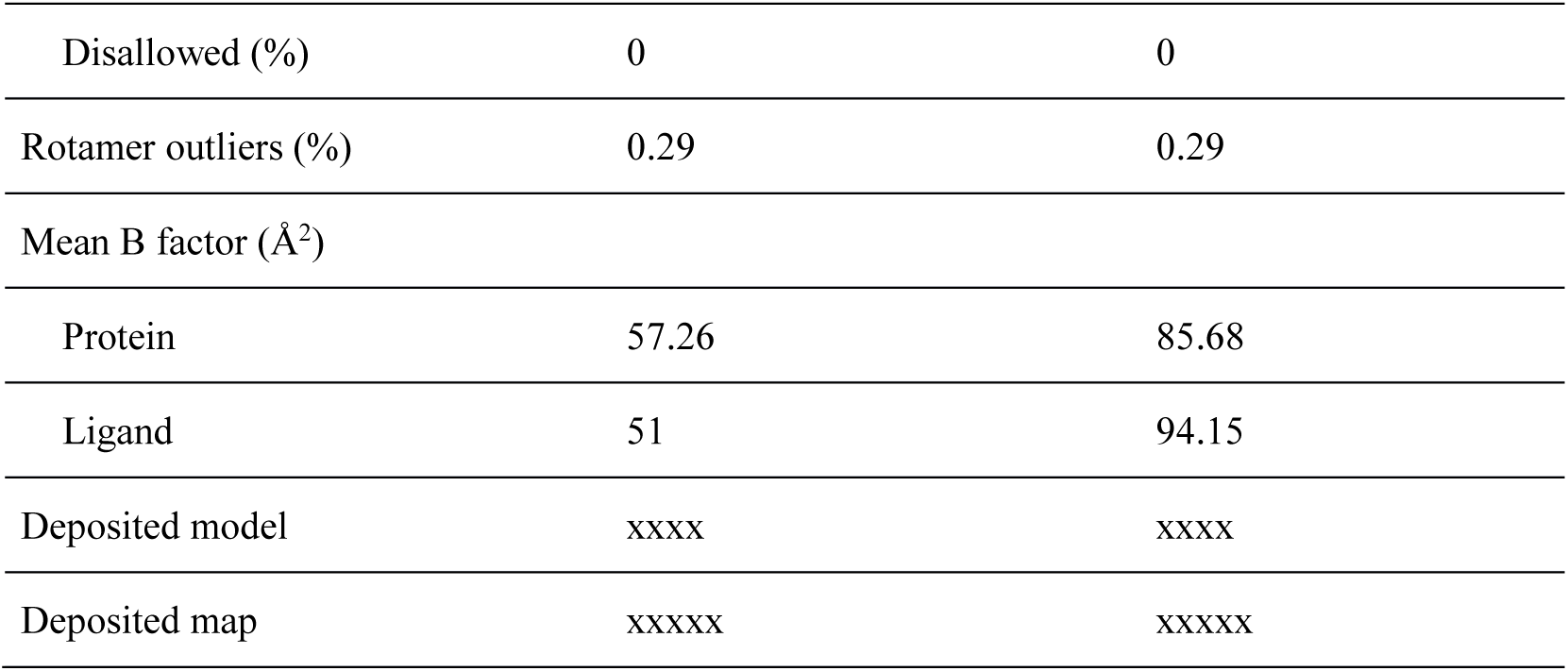

